# Aggregation of *Vibrio cholerae* by cationic polymers enhances quorum sensing but over-rides biofilm dissipation in response to autoinduction

**DOI:** 10.1101/333823

**Authors:** Nicolas Perez-Soto, Oliver Creese, Francisco Fernandez-Trillo, Anne-Marie Krachler

## Abstract

*Vibrio cholerae* is a Gram-negative bacterium found in aquatic environments and a human pathogen of global significance. Its transition between host-associated and environmental life styles involves the tight regulation of niche-specific phenotypes such as motility, biofilm formation and virulence. *V. cholerae*’s transition from the host to environmental dispersal usually involves suppression of virulence and dispersion of biofilm communities. In contrast to this naturally occurring transition, bacterial aggregation by cationic polymers triggers a unique response, which is to suppress virulence gene expression while also triggering biofilm formation by *V. cholerae*, an artificial combination of traits that is potentially very useful to bind and neutralize the pathogen from contaminated water. Here, we set out to uncover the mechanistic basis of this polymer-triggered bacterial behavior. We found that bacteria-polymer aggregates undergo rapid autoinduction and achieve quorum sensing at bacterial densities far below those required for autoinduction in the absence of polymers. We demonstrate this induction of quorum sensing is due both to a rapid formation of autoinducer gradients and local enhancement of autoinducer concentrations within bacterial clusters, as well as the stimulation of CAI-1 and AI-2 production by aggregated bacteria. We further found that polymers cause an induction of the biofilm specific regulator VpsR and the biofilm structural protein RbmA, bypassing the usual suppression of biofilm during autoinduction. Overall, this study highlights that synthetic materials can be used to cross-wire natural bacterial responses to achieve a combination of phenotypes with potentially useful applications.

## Introduction

Both natural and synthetic cationic macromolecules, such as cationic antimicrobial peptides, cationic polymers and dendrimers, have been extensively reported as antimicrobial.^1^ Due to their positive charge, these polymers can efficiently bind the negatively charged envelope of Gram-negative and Gram-positive bacteria.^2^ At high concentrations and charge densities, these molecules have the potential to interfere with membrane integrity and decrease bacterial viability.^1^ However, antimicrobial activity is heavily dependent on the length and nature of the polymer and, more importantly, on the nature of the targeted microbe. At low concentrations, cationic polymers are still capable of causing bacterial aggregation by mediating electrostatic interactions, but do so without significantly affecting bacterial membrane integrity and growth.^3^

We and others have previously reported that bacteria clustered by sub-inhibitory concentrations of cationic polymers display interesting phenotypes resembling those of biofilm communities.^3^ For instance, we have recently demonstrated that clustering of the diarrheal pathogen *Vibrio cholerae* with methacrylamides containing primary or tertiary amines leads to accumulation of biomass and extracellular DNA, and represses ToxT-regulated virulence factors including cholera toxin.^3f^ However, what drives these phenotypic changes in response to polymer exposure remains unclear. Similarly, the marine bacterium *Vibrio harveyi* shows enhanced bioluminescence in response to polymer-mediated clustering.^3a, 3b, 3e^ The bacterial components necessary to produce luminescence are encoded by the *luxCDABE* genes and subject to complex regulatory mechanisms. A major regulatory cascade controlling luminescence is quorum sensing. Since the regulators controlling luminescence are functionally conserved between *Vibrio* species, expression of *V. harveyi luxCDABE* genes can be used as a tool to probe quorum sensing in heterologous hosts.^4^

In *V. cholerae*, four parallel quorum sensing pathways operate,^5^ each governed by a autoinducer synthase, which produces a small, freely diffusible molecule that is released in the environment and can be sensed by a corresponding sensor/kinase component that controls the activity of the LuxU/LuxO phosphorelay. Of the four pathways, the LuxS/LuxPQ system, which produces and detects the inter-species autoinducer AI-2 (S-TMHF-borate) and the CqsA/CqsS system, which produces and senses the *Vibrio* specific autoinducer CAI-1 (S-3-hydroxytridecan-4-one), are the best characterized. While CqsA and LuxS synthesize the autoinducers, the hybrid sensor/kinases CqsS and LuxQ sense and respond to their cognate autoinducers by dephosphorylating and inactivating the regulator LuxO. When extracellular autoinducer concentrations are low, LuxO is phosphorylated and activates the transcription of the small RNAs Qrr1-4, which in turn inhibit the expression of HapR, a master regulator controlling diverse cellular functions including luminescence, biofilm formation and virulence.^6^ When the extracellular autoinducer concentration increases above a threshold, the sensor/kinases instead act as phosphatases, leading to dephosphorylation and inactivation of LuxO, ultimately allowing the expression of HapR. Consequently, when autoinducer concentration is high, HapR activates luminescence, but suppresses virulence and biofilm genes. This regulatory mechanism is thought to mediate the dissipation of host-associated biofilms and enable transmission of *V. cholerae* from the intestine to the environment.^7^ Since in *V. harveyi*, exposure to sub-inhibitory concentrations of cationic polymers lead to enhanced luminescence, here we set out to study whether *V. cholerae* would also activate quorum sensing in response to polymer-mediated clustering, and how quorum sensing could be related to the observed phenotypic changes, including lowered virulence and enhanced biofilm formation.

## Results

### Polymers enhance quorum sensing in *V. cholerae*

First, we set out to test if clustering by the cationic polymers poly(*N*-[3-aminopropyl] methacrylamide), (P1, Figure 1A) and poly(*N*- [3-(dimethylamino)propyl] methacrylamide), (P2, Figure 1B) induced quorum sensing in *V. cholerae*. As previously shown and detailed here by N-SIM super-resolution microscopy, even low concentrations of polymers induced rapid clustering of bacteria without affecting bacterial viability (Figure 1C). In order to create a fast and direct read-out for quorum sensing, the cosmid pBB1, which contains the *luxCDABE* luminescence genes from *V. harveyi,*^8^ was used to transform *V. cholerae* serogroup O1 biovar El Tor strain A1552 (serotype Inaba). The transformed bacteria were then grown in LB for 20 hours, washed with artificial marine water (AMW) and adjusted to an OD of 0.2 in AMW alone, or AMW containing polymers at concentrations ranging from 0.005 to 0.5 mg/ml. In the absence of polymer, luminescence as a read-out of quorum sensing first decreased due to back-dilution of the culture from a high density overnight culture to OD 0.2, and then gradually increased due to accumulation of autoinducers, peaking at 4.5 hours (Figure 1D and E, black traces). Interestingly, the quorum induction kinetics were significantly different in the presence of either P1 (Figure 1D) or P2 (Figure 1E), with initial luminescence being sustained and reaching a higher second peak at around 2.5 hours, as opposed to 4.5 hours in the absence of polymers. The magnitude of induction was higher in polymer-treated cultures (approx. 5-fold at peak quorum induction) compared to untreated cultures. Results were similar for the Ogawa serotype strain E7646, in that quorum sensing was initially sustained, and then further enhanced (approx. 4-fold at peak quorum induction) by polymer-mediated bacterial aggregation (Figure 1F, G). In both strains, the magnitude of luminescence induction was independent of the polymer concentration used, suggesting a threshold response.

**Figure 1.**
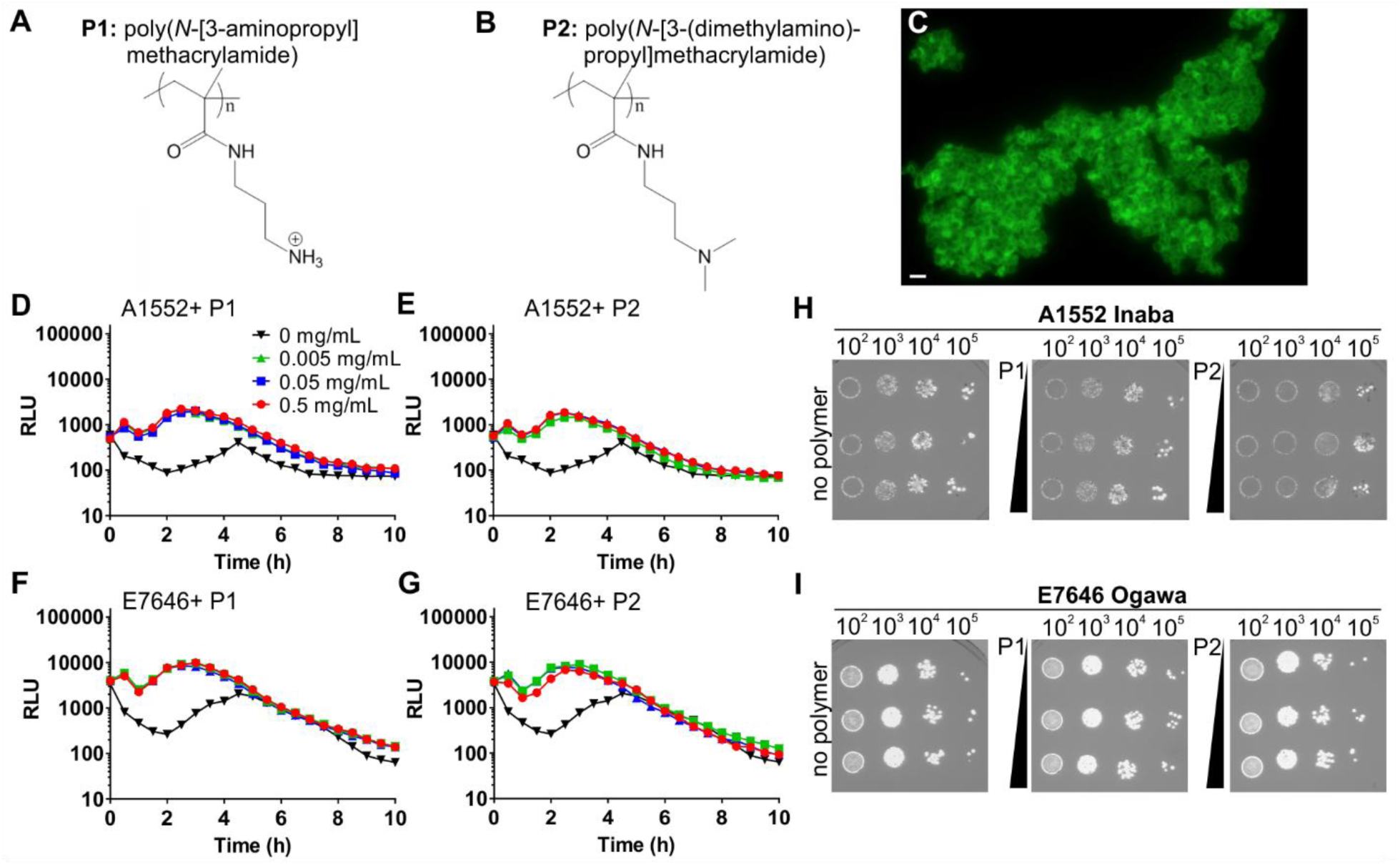
Polymers enhance quorum sensing in *Vibrio cholerae*. Chemical structures of P1 (A) and P2 (B). N-SIM super-resolution image of LIVE/DEAD-stained *V. cholerae* A1552 clustered by P1 (C). *V. cholerae* El Tor strains A1552 (D, E, H) or E7646 (F, G, I) containing the luminescence reporter pBB1 were adjusted to an OD_600_ of 0.2 following 16 hrs of growth and incubated in AMW alone (black) or AMW containing polymer P1 (D, F) or P2 (E, G) at concentrations of 0.005 (green), 0.05 (blue), or 0.5 (red) mg/ml. Luminescence was recorded every 30 min for 10 hrs and plotted as means ± s.e.m from at least three biological replicates. To test the effect of polymers on bacterial viability, samples were removed after 10 hrs, serially diluted and plated on LB (H, I).

To assess viable bacterial counts at the experimental end point, samples were serially diluted in high-salt buffer to disrupt clusters as previously described,^3f^ and plated. Similar numbers of colony forming units were recovered from untreated or polymer-treated samples (Figure 1H, I), suggesting that the presence of polymers did not affect bacterial viability or proliferation, in agreement with our previous data.^3f^ Taken together, our data demonstrate that these cationic polymers that cluster *V. cholerae* modulate the community behavior to give an accelerated and more robust autoinduction.

### Bacterial density shapes the kinetics of quorum induction in response to polymer

Since quorum sensing is usually tightly linked to bacterial density, we set out to explore how initial culture density affects quorum induction in the presence of polymers. *V. cholerae* carrying pBB1 as a quorum sensing reporter was adjusted to optical densities of 0.005, 0.05 and 0.5 in AMW alone, or AMW containing 0.005-0.5 mg/ml P1 or P2, and luminescence was monitored (Figure 2). The onset of autoinduction was not significantly modulated by the addition of polymers P1 or P2 to higher density cultures (OD_600_ of 0.5), but a ~ six-fold (P1) to nine-fold (P2) increase in peak luminescence was observed (Figure 2A, D). At lower culture densities, autoinduction of the untreated cultures was less pronounced (0.05) and eventually ceased (OD 0.005) since not enough autoinducer accumulated to reach the threshold concentration. In the presence of polymers, the initial quorum present in the culture was sustained, even in very dilute cultures, and the enhancement in peak luminescence was much more pronounced compared to untreated cultures (Figure 2B-F). Interestingly, clustering of very dilute cultures led to two peaks in luminescence, with a gap of approximately 4 hours (Figure 2C, F). Of note, the luminescence was not a result of bacterial growth, which was negligible under the observed conditions (AMW) and over the time frame described, both at high and low initial densities (Figure 2G, H).

**Figure 2.**
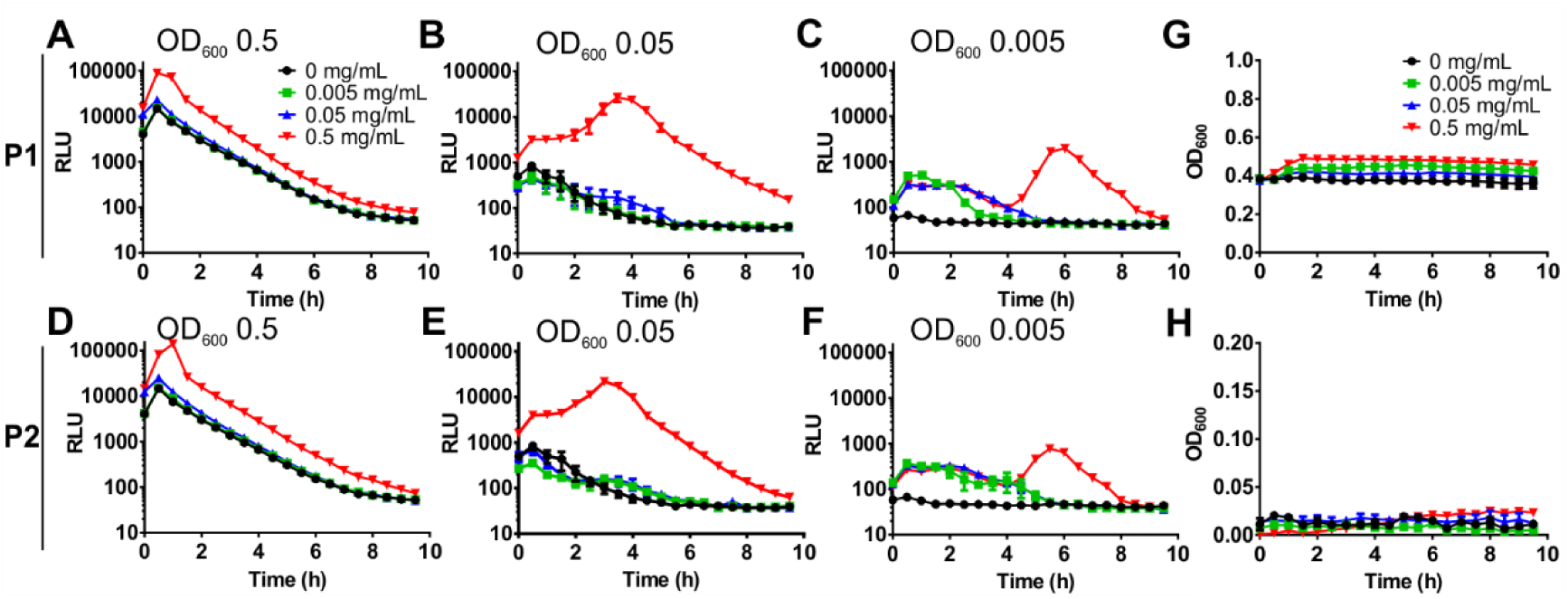
Bacterial density shapes the kinetics of quorum induction in response to polymer. Cultures of *V. cholerae* A1552 containing pBB1 were grown for 16 hrs and diluted into AMW alone (black) or AMW containing 0.005 (green), 0.05 (blue) or 0.5 (red) mg/ml of P1 (A-C) or P2 (D-F). Bacterial densities were adjusted to result in OD_600_ values of 0.5 (A, D), 0.05 (B, E), and 0.005 (C, F), respectively. Luminescence and OD_600_ were recorded every 30 min for 10 hrs and values are means ± s.e.m from at least three biological replicates. No significant growth was detected over this time frame, either at initial densities of 0.5 (G) or 0.005 (H).

To visualize the process of bacterial clustering and luminescence induction, *V. cholerae* A1552 containing pBB1 was incubated in the presence of P1, P2, or AMW alone in glass-bottom plates. Bacteria were imaged every 30 minutes to simultaneously visualize clustering and luminescence induction. With both P1 and P2 bacterial clusters were observed within minutes and remained stable over the duration of the experiment (Figure 3). Luminescence appears to originate from and to be restricted to bacterial clusters. Quantification of luminescence based on integrated pixel intensities showed a polymer-mediated enhancement of autoinduction (Figure 3H), in agreement with spectroscopic data.

**Figure 3.**
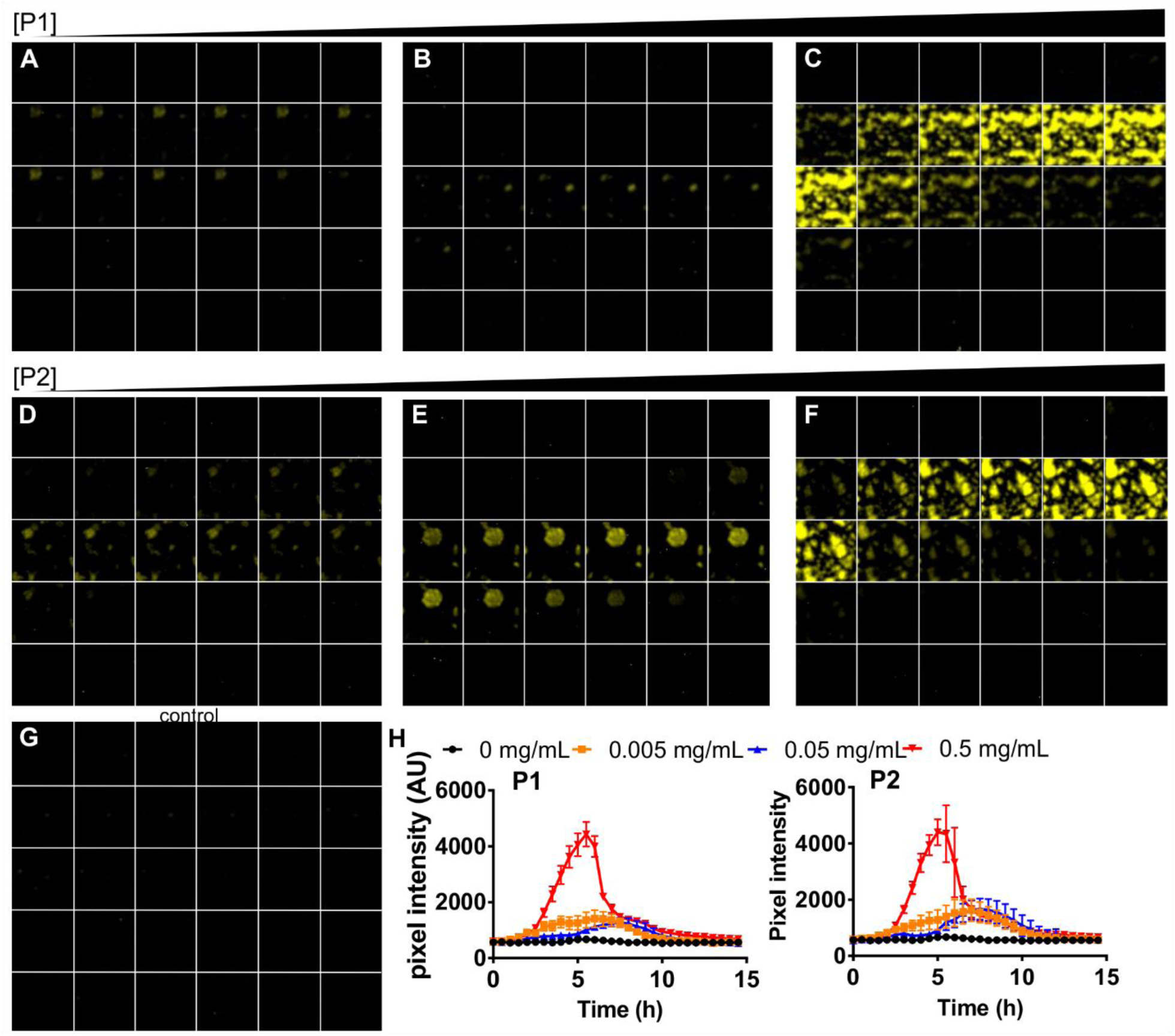
Polymers enhance quorum sensing in *Vibrio cholerae*. *V. cholerae* A1552 containing pBB1 was grown for 16 hrs and then adjusted to an OD600 of 0.2 either in the presence of P1 (A-C), P2 (D-F) or AMW alone (G). Polymers were adjusted to final concentrations of 0.005 (A, D), 0.05 (B, E), or 0.5 (C, F) mg/ml in AMW. Luminescence of samples was imaged every 30 min for 15 hrs and representative images for each time point are shown (0 hrs, top left to 15 hrs (bottom right) of each panel. Luminescence intensities over time were analyzed by quantifying pixel intensities, and means ± s.e.m from at least three biological replicates are shown for P1 (left) and P2 (right panel), (H).

### Polymer-mediated luminescence is not due to nutrient starvation within clusters

While autoinduction controls HapR and luminescence via the regulator LuxO, other environmental cues, including nutrient availability, have been reported to feed into the LuxO signaling cascade and thus have the potential to affect luminescence. Nutrient sensing and the LuxO signaling pathway converge at the cyclic AMP (cAMP) receptor protein, CRP.^9^ During limitation of PTS sugars such as glucose, cAMP-CRP is capable of modulating LuxO activation by affecting the expression of autoinducer synthases.^10^ Since clustering of bacteria by polymers may limit nutrient access and induce starvation, we tested whether a CRP deletion strain would be capable of activating luminescence in the presence of polymers. The *V. cholerae* Δ*crp* strain was transformed with pBB1 to monitor luminescence. However, the culture produced very low levels of luminescence, both in the presence and absence of polymers (Figure 4A, B), which is in agreement with previous work on *V. fischeri* CRP.^11^ While this suggests that cross-talk between CRP and LuxO signaling is a dominant cue for luminescence induction in *V. cholerae* as well, this made it unfeasible to determine whether the presence of polymers induced a state of carbon starvation, leading to luminescence induction via CRP. Instead, we repeated the experiment using the pBB1 containing *V. cholerae* wild type strain and supplementing the AMW with additional glucose. If clustering would limit nutrient diffusion, increasing the nutrient concentration should be able to overcome this and revert the bacteria to a non-luminescent phenotype. However, polymers still enhanced and sustained luminescence in the presence of 1% glucose, to a similar extent as in AMW alone (Figure 4C), suggesting that the effect was not due to nutrient limitation in the clusters.

**Figure 4.**
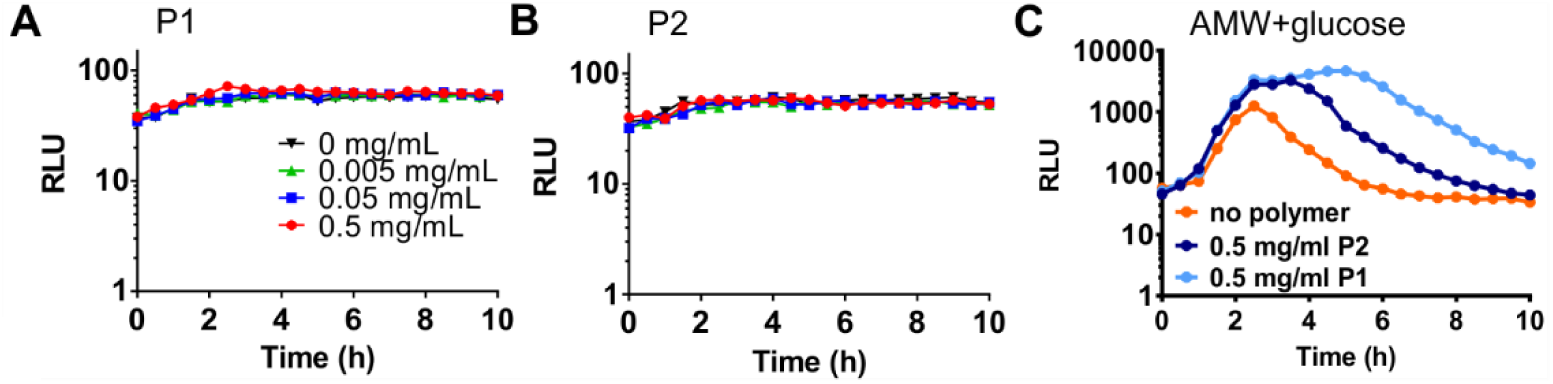
Polymer-mediated luminescence is not due to nutrient starvation within clusters. *V. cholerae* E7646 Δ*crp* containing the luminescence reporter pBB1 was grown for 16 hrs and then diluted to an OD_600_ of 0.2 in AMW alone (black) or AMW containing P1 (A) or P2 (B) at concentrations of 0.005 (green), 0.05 (blue) or 0.5 (red) mg/ml. Luminescence was recorded every 30 min for 10 hrs and means ± s.e.m from at least three biological replicates are shown. (C) *V. cholerae* E7646 wild type containing pBB1 was grown for 16 hrs and adjusted to an OD_600_ of 0.2 in AMW containing 1% glucose alone (orange), or in the presence of 0.5 mg/ml P1 (light blue) or P2 (dark blue). Luminescence was recorded every 30 min for 10 hrs and means ± s.e.m from at least three biological replicates are shown.

### Polymer mediated enhancement of quorum sensing is dominated by CAI-1

In *V. cholerae*, at least four parallel quorum sensing pathways converge to control the activity of the quorum sensing regulator, LuxO and thus, quorum regulated phenotypes including biofilm formation and virulence.^5^ We set out to test whether the polymer mediated effect on quorum sensing was specific for any one pathway. *V. cholerae* wild type strain A1552 was mixed with equal numbers of cells of quorum sensing mutants transformed with pBB1 either deficient in the production of AI-2 (DH231), or CAI-1 (WN1103), to give a final OD_600_ of 0.2. Additionally, the mutants were unable to sense the presence of AI-2 or CAI-1, respectively. Cultures producing less AI-2 produced a luminescence profile similar in shape and magnitude to quorum sensing proficient cells in response to polymer P1 addition (Figure 5A). In contrast, cultures producing less CAI-1 still had a comparable response profile upon addition of P1, but the magnitude of the luminescence enhancement was decreased (Figure 5B)_compared to cultures producing less AI-2 (Figure 5A). In co-cultures producing less of both autoinducers (containing strain BH1578), the luminescence response to polymer was abolished (Figure 5C). Similarly, co-cultures containing a low-density locked mutant of the down-stream quorum regulator LuxO (LuxO^D47E^) showed no luminescence induction upon addition of polymer P1 (Figure 5D). These data pointed at both autoinducers being involved in the quorum sensing enhancement in response to polymer, with CAI-1 being the dominant inducer. Additionally, the luminescence response to the polymer seems to proceed through the canonical LuxO-dependent pathway.

### Enhanced quorum sensing is driven by enhanced production of autoinducers in response to polymers

It has been described that at least under some conditions, quorum sensing may be subject to positive feedback, where quorum induction leads to increased production of one of the autoinducer synthases.^12^ Therefore, the enhancement in luminescence in response to polymers could be due to positive feedback, as a result of the polymers increasing the local concentration of autoinducers above the threshold. Alternatively, the polymers could have a direct effect on the production of autoinducers. We set out to test this by establishing a reporter assay that allowed us to decouple quorum sensing from the production of autoinducers. For this assay, we used as a luminescence reporter a *V. cholerae* strain transformed with pBB1 that could sense both CAI-1 and AI-2, but could not produce either molecule (BH1578). This reporter strain was exposed to supernatants from producer strains grown under different conditions, to evaluate the effect of the polymer. Initially, we evaluated the assay by growing the reporter in the presence of supernatants harvested from wild type *V. cholerae*, or strains incapable of producing either AI-2, CAI-1, or both autoinducers. Supernatant harvested from the quorum proficient wild type strain grown to high cell density triggered the highest level of luminescence in BH1578 (Figure 6A). The luminescence triggered by the AI-2 deficient strain was slightly decreased (Figure 6B), whereas luminescence was significantly decreased in response to supernatant from the CAI-1 deficient strain (Figure 6C) and was abolished in response to the strain deficient in both CAI-1 and AI-2 production (Figure 6D). Hence, the assay was capable of detecting different levels of autoinducers produced by a second strain.

We took this assay forward and harvested supernatants from wild type cells exposed to AMW alone, or AMW containing 0.005-0.5 mg/ml polymers P1 or P2, and exposed the reporter strain to filtered supernatants to test if the levels of autoinducers produced by the wild type strain in the presence of polymers were different. The reporter strain was not clustered under the assay conditions. While wild type cells exposed to AMW alone did not produce a detectable amount of autoinducer and thus, no significant luminescence reading in the reporter strain, both polymers P1 and P2 enhanced the production of autoinducers by wild type *V. cholerae*, leading to an increase in luminescence upon exposure of the reporter to the supernatants (Figure 6E, F). These data demonstrate that polymer exposure leads to enhanced production of autoinducers by the bacteria.

**Figure 5.**
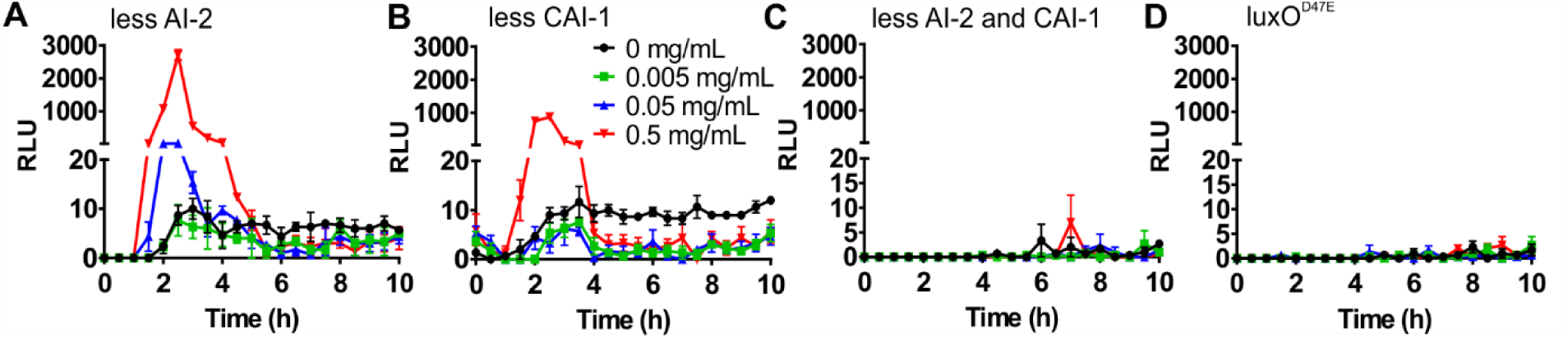
Polymer mediated enhancement of quorum sensing is mainly driven by CAI-1. *V. cholerae* A1552 wild type (dark) and quorum sensing mutants containing pBB1 were grown for 16 hrs, and then diluted into AMW to give equal cell densities and a total OD_600_ of 0.2. Strains were grown together in AMW alone (black) or AMW containing P1 at 0.005 (green), 0.05 (blue) or 0.5 (red) mg/ml. Luminescence was recorded every 30 min for 10 hrs and means ± s.e.m from at least three biological replicates are shown. Mutants grown in co-culture with the wild type were (A) DH231 (Δ*luxS*Δ*cqsS*) producing no AI-2, (B) WN1103 (Δ*luxQ*Δ*cqsA*) producing no CAI-1, (C) BH1578 (Δ*luxS*Δ*cqsA*) producing no AI-2 or CAI-1, and (D) BH1651 (luxO^D^47^E^).

**Figure 6.**
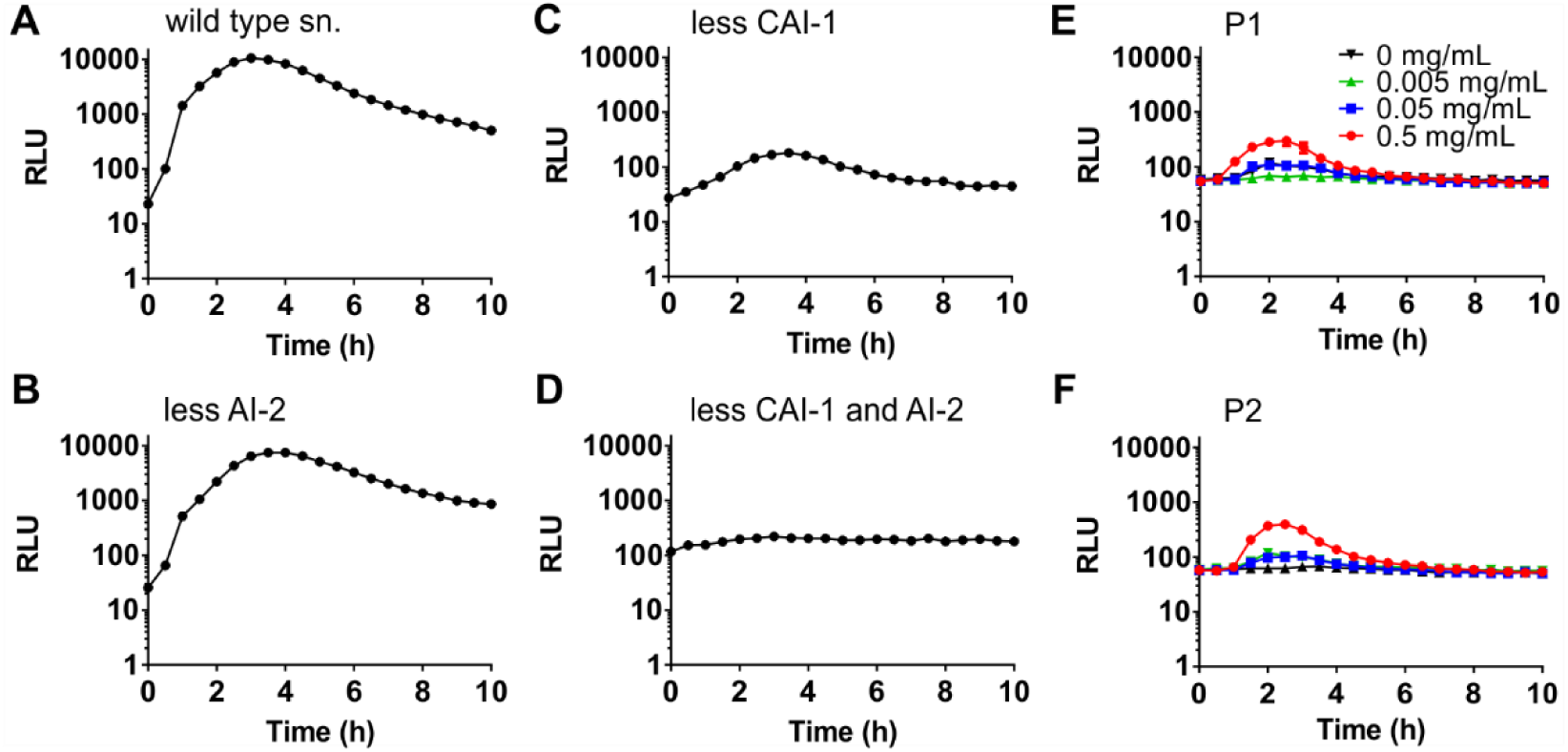
Enhanced quorum sensing is driven by enhanced production of autoinducers in response to polymers. Cultures of *V. cholerae* were adjusted to an OD of 0.2, grown for 16 hrs in LB medium, and supernatants were harvested, filtered, and incubated with *V. cholerae* BH1578 containing pBB1. Strains used to harvest supernatants were (A) wild type A1552 (B) DH231 (Δ*luxS*Δ*cqs*), (C) WN1103 (Δ*luxQ*Δ*cqsA*), and (D) BH1578 ((Δ*luxS*Δ*cqsA*). Luminescence was recorded every 30 min for 10 hrs and means ± s.e.m from at least three biological replicates are shown. *V. cholerae* wild type was adjusted to an OD_600_ of 0.2 in AMW alone or AMW containing 0.005-0.5 mg/ml P1 (E) or P2 (F) and supernatants harvested and filtered 16 hrs later. To determine their autoinducer content, supernatants were incubated with the reporter strain *V. cholerae* BH1578 containing pBB1. Luminescence was recorded every 30 min for 10 hrs and means ± s.e.m from at least three biological replicates are shown.

### Polymer-mediated quorum induction over-rides the canonical biofilm dissipation programme in *V. cholerae*

Contrary to many bacteria that use quorum signaling as a means to induce biofilm formation, in *V. cholerae* autoinduction promotes repression of biofilm production and dissemination, via the regulator HapR.^13^ However, we had previously observed enhanced accumulation of *V. cholerae* upon prolonged exposure to cationic polymers, but whether this was accompanied by transcriptional changes at the level of biofilm production was not known. Thus, we grew *V. cholerae* containing transcriptional fusions to promoters regulating key biofilm components in the presence or absence of P1 and P2 (Figure 7). *V. cholerae* biofilms contain the structural protein RbmA and require the regulator VpsR, which controls the expression of the *vps* polysaccharide biosynthesis genes.^14^ Upon exposure to either P1 or P2, *vpsR* and *rbmA* were both significantly induced (Figure 7A, B), suggesting that upon polymer-mediated clustering, quorum sensing does not, as usually, suppress genes involved in biofilm production, but instead their transcription is enhanced. AphA, which is another direct target and is usually induced by VpsR, is suppressed in the presence of polymers (Figure 7C).

**Figure 7.**
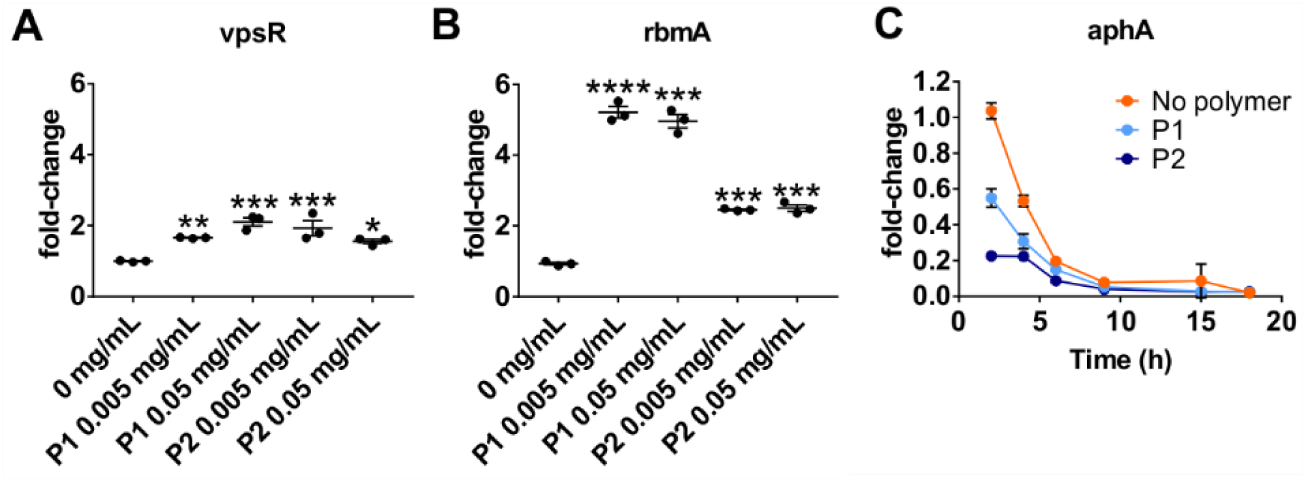
Polymer-mediated quorum induction over-rides the canonical biofilm dissipation programme in *V. cholerae*. *V. cholerae* wild type A1552 containing pRW50T lacZ reporters for the promoters of *rbmA* (A), or *vpsR* (B), were grown for 16 hrs and then diluted into AMW alone or containing 0.005 or 0.05 mg/ml P1 or P2 as indicated, to give an OD_600_ of 0.2. Following 16 hrs of incubation, clustered bacteria were removed and either processed for beta-galactosidase assays or treated with high-salt PBS to disperse the cultures for OD_600_ measurements. Transcriptional activities were calculated and normalized to untreated cultures. Shown are means ± s.e.m and individual measurements for three biological replicates. Statistical significance was determined by ANOVA and a Dunnett’s multiple comparison test and is depicted as (****) for p-values ≤ 0.0001, (***) p≤0.001, (**) p≤0.01 (*) p≤0.05 and ns or not significant (p≥0.05). (C) *V. cholerae* wild type A1552 containing pRW50T lacZ reporter for the *aphA* promoter was grown in AMW alone (orange) or containing P1 (light blue) at 0.05 mg/mL or P2 (dark blue) at 0.5 mg/mL for 18 hrs. Clustered bacteria were removed at indicated times and either processed for beta-galactosidase assays or treated with high-salt PBS to disperse the cultures for OD_600_ measurements. Transcriptional activities were calculated and normalized to the activities of untreated cultures at 2hrs. Shown are means ± s.e.m and individual measurements for three biological replicates.

## Discussion

Traditionally, work on cationic polymers has been carried out with the development of antimicrobial materials as a main goal.^1^ However, recent work by our groups and others has demonstrated that such cationic polymers can be titrated against bacteria, to achieve a charge balance that allows for the rapid and efficient clustering of bacteria but avoids membrane disruption and bacterial cell death.^3^

The use of cationic polymers to induce rapid bacterial clustering in this way has proven as an interesting path to study effects of cell aggregation and crowding on bacterial physiology. While such behaviors are often studied in batch cultures, by incubating bacterial cultures over a prolonged time, this means aggregation is accompanied by bacterial growth, and eventually nutrient limitation, which makes it difficult to establish the primary cause of the observed phenotypes. In contrast, cationic polymers induce cell aggregation rapidly, within minutes, which allows us to study these phenomena independent of cellular proliferation and nutrient limitation.

We and others have previously observed that cationic polymers and dendrimers can, under certain conditions, trigger bioluminescence in the marine bacterium *V. harveyi*, suggesting they may induce or enhance quorum sensing.^3a, 3b, 3d-3e^ In a more recent study where we extended this work to the human pathogen *V. cholerae*, we observed that polymer-mediated clustering led to enhanced deposition of biomass and extracellular DNA, while it interfered with the induction of virulence genes in an infection model.^3f^ Since virulence and biofilm production are both regulated by quorum sensing but are usually both regulated concurrently, the goal of this study was to test whether cationic polymers would trigger quorum sensing in *V. cholerae* and how this would affect down-stream transcription of biofilm genes.

We used *V. cholerae* strains heterologously expressing the *luxCDABE* luminescence genes (on cosmid pBB1) from *V. harveyi* to be able to use luminescence as direct readout for autoinduction. Over 16 hours, *V. cholerae* would grow to high cell densities and as a result, was strongly luminescent. On dilution into artificial marine water, cell density and autoinducer concentration would rapidly decrease, resulting in a decline in luminescence. After several hours, cells would eventually accumulate sufficient autoinducer to reach the quorum threshold and induce luminescence again. This behavior was observed in AMW alone (Figure 1) and is in agreement with commonly observed results from such experiments.^5, 12^ In contrast, when cells were diluted into media containing polymers, they would undergo extensive clustering almost instantaneously, and luminescence readouts never dropped, but instead, further increased immediately (Figure 1), suggesting that clustering not only countered the dilution effect, but further increased autoinducer concentration within the clusters. Interestingly, this behavior was observed over a broad space of cell densities (at least two orders of magnitude), including in dilute cultures that did not by themselves experience autoinduction (Figure 2C, F), suggesting that during clustering, polymers create pockets containing strongly increased concentrations of autoinducers around bacterial aggregates.

We further demonstrated that both CAI-1 and AI-2 dependent quorum sensing cascades are activated in response to polymers and that clustering leads to an enhanced production of both autoinducers (Figures 5 and 6). The effect of the *Vibrio*-specific autoinducer CAI-1 dominated the clustering-driven luminescence phenotype (Figure 5), in line with previous results obtained for batch-cultures of *V. cholerae* in rich medium.^12^

Some studies have hypothesized that luminescence could be a result of limited diffusion of nutrients in the polymer-mediated bacterial aggregates.^3d^ Catabolite repression of luminescence has been reported for *V. fischeri*, where cAMP-CRP stimulates *luxCDABE* expression.^15^ However, this effect is alleviated by high concentrations of autoinducer.^11^ In our hands, CRP was essential for luminescence, both triggered by high cell density in the absence of polymers, in line with previous findings for an *E. coli* Δ*crp* mutant,^11^ as well as in response to polymer-induced clustering. Additionally, supplementation of the media with excess glucose did not quench luminescence, even in the absence of autoinduction. This suggests that nutrient limitation within the clusters is not a major cue for luminescence induction, but further underpins that cross-talk between nutrient sensing and quorum sensing pathways exists.

Finally, we followed up on our earlier observation that exposure to cationic polymers causes deposition of *V. cholerae* on inorganic surfaces and release of extracellular DNA, both hallmarks of biofilm formation.^3f^ Here, we showed that this phenotype is the result of transcriptional activation of genes involved in biofilm production in response to polymer exposure. Biofilm induction may explain the enhanced resistance towards antimicrobials of bacteria that have been exposed to cationic polymers, as previously described by others.^3d^ The expression of the biofilm regulator VpsR and the biofilm structural protein RbmA were both induced upon exposure to the polymers (Figure 7). This upregulation is in contrast to the canonical biofilm regulation where biofilm genes are repressed during autoinduction. VpsR is a master regulator of biofilm formation and a two component system response regulator. Although no cognate histidine kinase has been identified, VpsR is epistatic to the intracellular hybrid sensor histidine kinase VpsS.^16^ Induction of *vpsR* likely leads to the downstream induction of *rbmA* we observed here, since *rbmA* is a direct target of VpsR regulation.^17^ However, *vpsR* induction in the presence of polymers seems to happen despite autoinduction, which should normally lead to suppression of *vpsR*. What is also different from a regular biofilm response is that VpsR in the presence of polymer, fails to upregulate one of its other direct targets, *aphA*. We showed that in contrast to this canonical response, *aphA* is strongly suppressed by the presence of polymers (Figure 7C). When Shikuma et al. identified VpsS as a regulator of VpsR, they established the existence of a pathway that proceeds from VpsS through the quorum regulators LuxU and LuxO and results in the VpsR dependent activation of biofilm production, independent of HapR.^16^ It may be that in the presence of polymers, this pathway is active and dominates the effects of the CAI-1 and AI-2 pathways on biofilm. Unfortunately, the cognate signal activating VpsS is as yet unidentified.

## Conclusions

We showed here that clustering of *V. cholerae* in response to cationic polymers leads to autoinduction, due to a rapid increase of local autoinducer concentration in the vicinity of aggregated bacteria. Moreover, we demonstrate that stimulation of further autoinducer synthesis is also observed and involves at least two of the four known quorum sensing systems, CAI-1 and AI-2. We speculate that the third quorum sensing pathway, which proceeds through the intracellular hybrid sensor kinase VpsS ^5, 16^ is also activated, and leads to the production of biofilm in response to polymer driven aggregation. Our previous work together with the data presented here rules out membrane disruption and nutrient limitation within clusters, respectively, as cues leading to the phenotypes observed here. Our future work will aim to further dissect the pathway(s) triggered in response to polymer exposure, to clarify whether VpsS is indeed involved, and is activated in response to polymers. Polymeric materials that inhibit bacterial dissemination, both mechanically and transcriptionally, may be useful for applications to enhance waste water treatment for decontamination of water from *V. cholerae*.

## Materials and Methods

### Bacterial strains and culture conditions

*V. cholerae* El Tor strains used in this study (Table 1) were derived from A1552 ^18^ and E7946.^19^ The *E. coli* K12 strains JCB387 ^20^ DH5α ^21^ and SM10 λpir ^22^ were used for general cloning and conjugation procedures. Strains were propagated at 37°C in lysogeny broth (LB) supplemented with 10 μg/μL tetracycline or 30 μg/μL kanamycin for selection when required. Plasmids were introduced into *V. cholerae* strains by triparental mating with *E. coli* DH5a carrying the desired plasmid (donor) and *E. coli* SM10 (helper strain) carrying the conjugative machinery on pRK2013. Cultures were mixed at a volumetric ratio of 1:2:2 of recipient:helper:donor in 250 µl and spotted onto brain-heart infusion (BHI) agar to be incubated overnight at 37 **°**C. Spots of bacteria were dislodged after an overnight incubation and resuspended in 3 mL of sterile PBS. 100 μL of serial dilutions were plated onto TCBS plates containing 10 μg/μL of tetracycline. Resulting colonies were checked by PCR in the case of pRW50T constructs, while pBB1 transconjugants were screened for luminescence.

**Table 1.** Bacterial strains used in this study.

| Strain | Description or genotype | Source or Reference |
| --- | --- | --- |
| *Vibrio cholerae* |  |  |
| A1552 | Wild-type; O1 biovar El Tor serotype Inaba | 18 |
| E7956 | Wild-type; O1 biovar El Tor serotype Ogawa | 19 |
| BH1651 | *luxO*<sup>D47E</sup> | 6 |
| BH1578 | $\Delta luxS \Delta cqsA$ | 25 |
| DH231 | $\Delta luxS \Delta cqsS$ | 26 |
| WN1103 | $\Delta luxQ \Delta cqsA$ | 26 |
| E7956 $\Delta crp$ | $\Delta crp$ Kan<sup>R</sup> | Gift from D. Grainger |
| NP5005 | A1552 pRW50T containing upstream region of *aphA* promoter; Tet<sup>R</sup> | 3f |
| *Escherichia coli* |  |  |
| DH5α | Donor and maintenance of pBB1 | 21 |
| JCB387 | Donor and maintenance of pRW50T | 20 |
| SM10 | Helper strain; $\lambda pir$ pRK2013; Kan<sup>R</sup> | 22 |

**Table 2.**
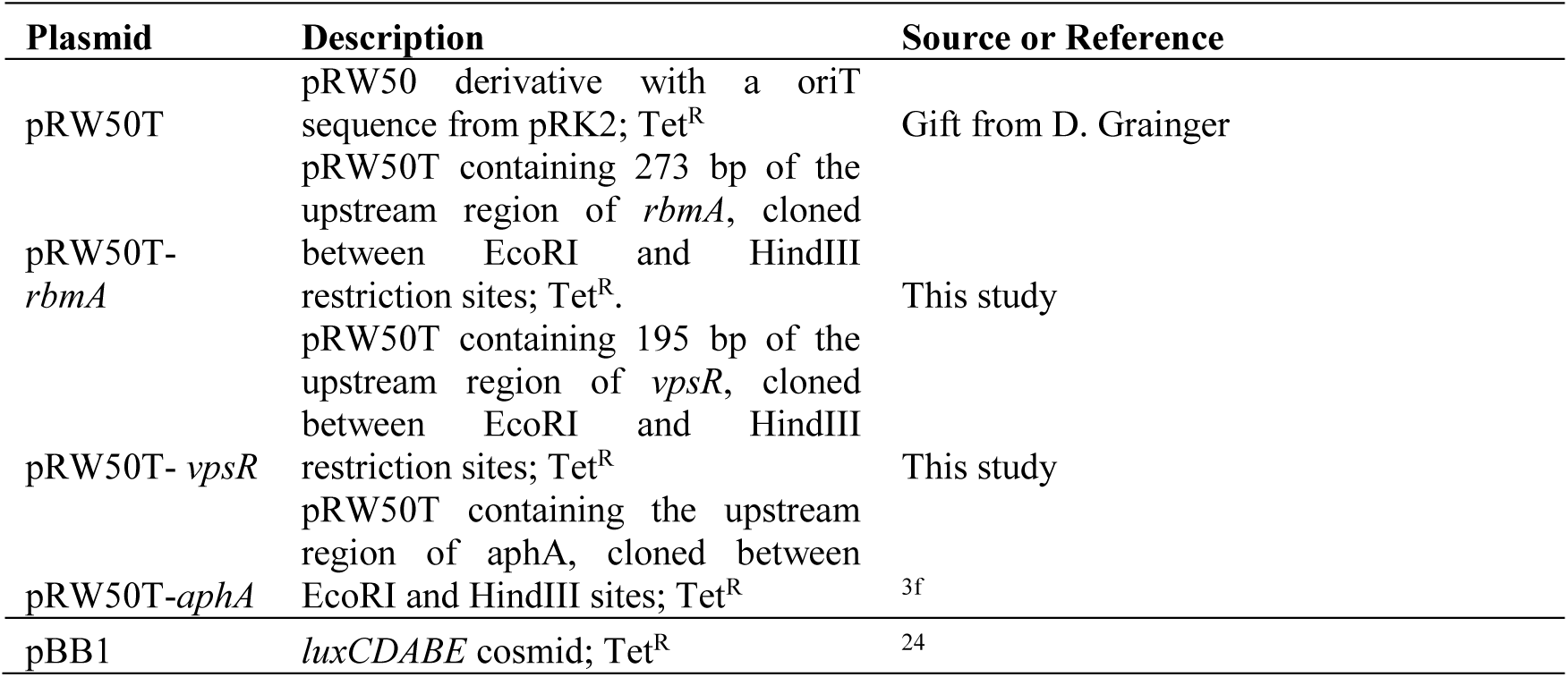
Plasmids used in this study.

### Beta-galactosidase assays

pRW50T derivative construction was described before.^3f^ Regions encoding *aphA, rbmA*, or *vpsR* promoters were amplified by PCR and cloned into pRW50T using EcoRI and HindIII sites. The insertion was checked by PCR using external primers. Measurement of β-galactosidase activity as a readout for transcriptional activity was done as previously described ^23^, with some modifications to accommodate testing of aggregated bacteria. Small cultures of reporter strains were grown in the absence or presence of polymers P1 and P2 and incubated overnight at 37 °C with shaking. Clustered bacteria were split in two, and either used for transcriptional assays or washed with high salt PBS (200 mM NaCl) to disrupt aggregation and enable OD_600_ measurements.

### Luminescence assays

Luminescence assays were done using *V. cholerae* pBB1 transconjugants. The pBB1 cosmid^24^ was introduced into *V. cholerae* strains by triparental mating in the same conditions as for pRW50T. Overnight cultures of *V. cholerae* pBB1 were adjusted to OD_600_ of 0.5, 0.1 and 0.01 in artificial marine water with 10 μg/μL of tetracycline. Polymers P1 and P2 were added at concentrations of 0.005 mg/mL, 0.05 mg/mL and 0.5 mg/mL in DMEM or AMW, in 200 μL final volume using a dark wall clear bottom 96-well plate. Plate was incubated up to 15 hours at 37°C with shaking a 200 rpm, while luminescence and OD_600_ were recorded every 30 minutes using a FLUOstar Omega plate reader. The following assays were done with bacterial cultures with OD_600_ adjusted to 0.2. Cells were recovered after the assay and washed with high-salt PBS containing 200 mM NaCl to disrupt charge-based aggregation, and plated onto LB with tetracycline to determine the viability. Plates were imaged using a BioRad Gel Doc XR System and images were processed with ImageJ.

### Luminescence time-lapse imaging

Overnight culture of *V. cholerae* A1552 pBB1 was diluted to an OD_600_ of 0.2 in artificial marine water or clear DMEM with 10 μg/μL of tetracycline, and polymers at concentrations of 0.005 mg/mL, 0.05 mg/mL and 0.5 mg/mL. Samples were prepared in 200 μL using a glass-bottom 96-well plate and incubated at 37 °C with 5% CO_2_ in a microscope imaging chamber. Images were taken every 30 minutes with 10 seconds of exposure at 40X magnification, using an Evolve 512 EMCCD camera mounted on a Nikon-Eclipse TE2000-U microscope. Image acquisition was done using Nikon NIS-Elements software and final images processed with ImageJ. Pixel intensity was determined from several clusters within frame using ImageJ.

### Super resolution microscopy of bacterial clusters

*V. cholerae* A1552 was incubated with 0.05 mg/ml P1 in PBS for 1 hour. To visualize membrane integrity, the sample was stained using the LIVE/DEAD BacLight Bacterial Viability Kit (Life Technologies) for 10 minutes at room temperature. The sample was mounted with ProLong Gold Antifade Mountant and covered with a cover slip. Images were taken on a Nikon N-SIM super resolution microscope fitted with SR Apo TIRF 100x lens, at 100 ms exposure. Deconvolution was carried out using the Nikon NIS elements software.

### Luminescence assays using *V. cholerae* BH1578 pBB1 as a reporter

*V. cholerae* BH1578 pBB1 was used to determine the effect of polymers on the production of autoinducers. *V. cholerae* strains at an OD_600_ of 0.2 were clustered with polymers at concentrations of 0.005 mg/mL, 0.05 mg/mL and 0.5 mg/mL in artificial marine water. Supernatants were recovered by centrifugation, and used to resuspend *V. cholerae* BH1578 pBB1 previously adjusted to an OD_600_ of 0.2 in 200 μL. Luminescence was recorded at 37 °C using a FLUOStar Omega plate reader. Similarly, *V. cholerae* strains and *V. cholerae* BH1578 pBB1 were co-cultured in 200 μL final volume and polymers added at concentrations of 0.005 mg/mL, 0.05 mg/mL and 0.5 mg/mL in artificial marine water. Both strains were adjusted to a final OD_600_ of 0.1 each (0.2 total density). Incubation was done at 37 °C with shaking at 200 rpm, and luminescence and OD_600_ were measured every 30 minutes using a FLUOStar Omega plate reader.

## Acknowledgements

We would like to thank B. Bassler for the generous gift of quorum sensing mutants BH1578, DH23, WN1103 and the cosmid pBB1. We would like to thank H. Kaplan, the Krachler and Fernandez-Trillo labs for critical reading of the manuscript, and for suggestions on how to improve this study. This work was supported by University of Birmingham Fellowships (to A.M.K. and F.F.-T.), Wellcome Trust grant 177ISSFPP (to A.M.K. and F.F.-T), BBSRC grants BB/M021513/1 (to A.M.K.) and BB/L007916/1 (to A.M.K.), a CONICYT fellowship (to N.P.- S.), BBSRC MIBTP scholarship BB/M01116X/1 (to O.C.) and a UT Systems Science and Technology Acquisition and Retention Award (to A.M.K.).

